# Toward a Monte Carlo approach to selecting climate variables in MaxEnt: A case study using Cassin’s Sparrow (*Peucaea cassinii*)

**DOI:** 10.1101/2020.07.15.202945

**Authors:** John L. Schnase, Mark L. Carroll, Roger L. Gill, Glenn S. Tamkin, Jian Li, Savannah L. Strong, Thomas P. Maxwell, Mary E. Aronne

## Abstract

MaxEnt is an important aid in understanding the influence of climate change on species distributions and abundance. There is growing interest in using IPCC-class global climate model outputs as environmental predictors in this work. These models provide realistic, global representations of the climate system, projections for hundreds of variables (including Essential Climate Variables), and combine observations from an array of satellite, airborne, and *in-situ* sensors. Unfortunately, direct use of this important class of data in MaxEnt modeling has been limited by the large size of climate model output collections and the fact that MaxEnt can only operate on a relatively small set of predictors stored in a computer’s main memory. In this study, we demonstrate the feasibility of a Monte Carlo method that overcomes this limitation by finding a useful subset of predictors in a larger, externally-stored collection of environmental variables in a reasonable amount of time. Our proposed solution takes an ensemble approach wherein many MaxEnt runs, each drawing on a small random subset of variables, converges on a global estimate of the top contributing subset of variables in the larger collection. In preliminary tests, the Monte Carlo approach selected a consistent set of top six variables within 540 runs, with the four most contributory variables of the top six accounting for approximately 93% of overall permutation importance in the final model. These results suggest that a Monte Carlo approach could offer a viable means of selecting environmental predictors for MaxEnt models that is amenable to parallelization and scalable to very large data sets. This point to the possibility of near-real-time multiprocessor implementations that could enable broader and more exploratory use of global climate model outputs in environmental niche modeling and aid in the discovery of viable predictors.

## Introduction

MaxEnt is one of the most popular software packages in use today by the ecological research community [1–3]. Based on a machine learning approach to maximum entropy modeling, MaxEnt allows researchers to construct ecological niche models (ENMs) that estimate the habitat suitability of a species using occurrence data and a set of environmental variables [1,2,4–6]. An abundant literature points to MaxEnt’s effectiveness across a wide range of applications in fields as diverse as biogeography and phylogeny [7], conservation biology and epidemiology [8,9], invasion biology [10–12], and archaeology [13]. Its merits compared to alternative approaches have been the subject of numerous statistical and methodological analyses, many of which have led to software improvements and refinements to the way MaxEnt is used [14–21]. In this paper, we contribute to this ongoing dialog by describing our efforts to overcome a specific technical limitation of the MaxEnt software that makes the tool difficult to use with large predictor data sets.

In recent years, MaxEnt has become a particularly important aid in understanding the influence of climate change on species distributions and abundance [22–27]. The need for reliable climate projections in this work is leading to greater use of global climate model (GCM) outputs as predictors [23]. While creating important new opportunities for research, this trend is also creating a “Big Data” challenge for the MaxEnt community [16]. The largest and most sophisticated GCMs — sometimes referred to as “IPCC-class” models because of the critical role they play in the work of the Intergovernmental Panel on Climate Change (IPCC) — produce petabyte-scale data sets comprising hundreds of variables, a volume that vastly exceeds what is generally used in bioclimatic modeling today [28–30]. Moreover, the direct outputs of these systems are being transformed into derived climate data products on an unprecedented scale [24,31]. As a result, model tuning and variable selection, which are crucial aspects of any species distribution modeling effort, are becoming more complicated issues [32].

Part of the problem lies in the fact that MaxEnt, like many machine learning systems, acts on its inputs as a piece: predictors and observations must be memory-resident for the program to work [33]. This results in run-times and space requirements that scale linearly with the size of a model’s inputs. In most cases, these scaling properties pose few difficulties. But when the number of predictors under consideration becomes large, compute times can become impractically long, models can become overly complex, and efforts to understand any particular variable’s contribution to model formation, either as an aspect of model analysis or as a way of selecting subsets of variables for further model refinement, can become challenging [17,32,34–36]. Clearly, an effective way of dealing with large environmental data sets that preserves the many advantages of MaxEnt while overcoming its current limitations would benefit the MaxEnt community.

In this study, we investigated the potential of a Monte Carlo method to help accomplish such an outcome. Monte Carlo optimizations are a common way of finding approximate answers to problems that are solvable in principle but lack a practical means of solution [37]. Our objective was to find a useful subset of predictors in a larger collection of environmental variables in a reasonable amount of time. Our proposed solution takes an ensemble approach wherein many MaxEnt runs, each drawing on a small random subset of variables, converges on a global estimate of the top contributing subset of variables in the larger collection.

Preliminary results suggest that the method reliably selects a subset of the original predictors that is capable of producing a well-tuned, parsimonious model of high quality. Since each model run is independent and uses a set number of variables, the method is totally parallelizable, independent of the scaling properties of MaxEnt, and amenable to implementation as an external memory algorithm. If proved to be effective, such an approach could provide a practical way of constructing MaxEnt models when there is a need to select a small set of predictors in a pool comprising a potentially very large number of predictors. This could lead to greater use of climate model outputs by the ecological research community and aid the search for viable predictors when variable selection through ecological reasoning is not apparent.

## Materials and Methods

Cassin’s Sparrow (*Peucaea cassinii* Woodhouse, 1852) is an elusive resident of arid shrub grasslands of Middle America and the Southwestern United States [38]. Desert-adapted birds, such as Cassin’s Sparrow, appear to be especially vulnerable to climate change [39,40]. We chose Cassin’s Sparrow as a target for our development efforts as an example of a species whose study could benefit from the technical advances described here. Occurrence data was obtained from the Global Biodiversity Information Facility (GBIF) for the year 2016 [41]. After removing replicates, a total of 1865 records were acquired. To limit spatial extent and avoid pseudo-replication, we thinned the points to a radius of 16 km, which resulted in a total of 609 observations. For predictors, we used Worldclim’s standard 19 Bioclimatic (bioclim) environmental variables at a resolution of 2.5 arc-minutes throughout (Table 1) [42,43]. These predictor layers were clipped to the coverage area of our observational data, reprojected, and formatted for use with MaxEnt using the Geospatial Data Abstraction Library (GDAL) software package [44] following the guidelines of Hijmans et al. [45]. We did not attempt to minimize collinearity by removing variables, because the current study focuses on an assessment of stochastic down-selection from a full variable set, and because MaxEnt has a demonstrated ability to account well for redundant variables [46].

**Table 1.** Worldclim Bioclimatic Variables.

|  |  |
| --- | --- |
| bio01 | Annual Mean Temperature |
| bio02 | Mean Diurnal Range (Mean of monthly (max temp - min temp)) |
| bio03 | Isothermality (BIO2/BIO7) ($\times 100$) |
| bio04 | Temperature Seasonality (standard deviation $\times 100$) |
| bio05 | Max Temperature of Warmest Month |
| bio06 | Min Temperature of Coldest Month |
| bio07 | Temperature Annual Range (BIO5-BIO6) |
| bio08 | Mean Temperature of Wettest Quarter |
| bio09 | Mean Temperature of Driest Quarter |
| bio10 | Mean Temperature of Warmest Quarter |
| bio11 | Mean Temperature of Coldest Quarter |
| bio12 | Annual Precipitation |
| bio13 | Precipitation of Wettest Month |
| bio14 | Precipitation of Driest Month |
| bio15 | Precipitation Seasonality (Coefficient of Variation) |
| bio16 | Precipitation of Wettest Quarter |
| bio17 | Precipitation of Driest Quarter |
| bio18 | Precipitation of Warmest Quarter |
| bio19 | Precipitation of Coldest Quarter |

We used MaxEnt Version 3.4.1 [47], R Version 4.0.1 [48], the ENMEval Version 0.3.0 R package [49], RStudio Version 1.2.5033 [50], and ENMTools Version 1.4.4 [51] running on a 2.8 GHz Intel Core i7 MacBook Pro with 16 GB of memory in the study. First, we developed a baseline model using the stand-alone MaxEnt program operated through its graphical user interface (GUI). MaxEnt users can apply various combinations of five mathematical transformations (‘feature classes’ or FCs) to predictor variables to enable more complex fits to the observational data. The available feature types for continuous variables are linear (L), quadratic (Q), hinge (H), product (P), and threshold (T) [4]. Users can also adjust a regularization multiplier (RM) to maximize predictive accuracy and offset the overfitting that FC adjustments can introduce. We applied MaxEnt’s default FC and RM settings (i.e. the “Auto features” setting) with 10 replicate cross-validation and jackknife evaluation of variable importance. By default, MaxEnt uses all feature classes and a regularization multiplier of 1.0 when there are more than 80 training samples, which was the case here [47]. Ten thousand background points were selected from across the study area following the recommendations of Phillips et al. [52] and Fourcade et al. [53]. We determined the average permutation importance for each variable in three replicated runs. The top six predictors in the three-run ensemble were used to develop the final MaxEnt baseline model.

We then developed an alternative method to select the top six variables that is based on random sampling. We implemented our Monte Carlo approach as an R script that invokes MaxEnt through ENMEval, which provides convenient control over model settings, built-in evaluation metrics, and improved performance [35,49]. To reduce variability and isolate outcomes as much as possible to the effects of the sampling process, we adopted a feature class setting of LQHP and a regularization multiplier setting of 1.0 as fixed parameters in all the Monte Carlo runs. We defined ensemble, in this case, to mean a collection of 100 sprints, where each sprint consisted of ten model runs. A tally table was used to maintain a count of the number of times a variable was used in a model run along with a cumulative sum of the variable’s permutation importance. The tally table thus provided the information needed to determine the average permutation importance of a predictor at any point along the way.

To process a sprint, we initialized each of its ten model runs with a random subset of environmental variables read from the filesystem. Random integers drawn from a uniform distribution ranging 1–19 corresponding to the 19 bioclim predictors were used to make the selection. At the conclusion of each model run, the tally table was updated appropriately. At the conclusion of each sprint, we computed a MaxEnt model using the six predictors in the original starting set having the highest average permutation importance values at that point. This process was repeated 100 times to produce a complete ensemble. We assessed the algorithm’s performance in two ensembles. In the first, we chose two random variables for each sprint run; in the second, six random variables were used for each run. This resulted in an overall total of 2000 model runs.

The predictive distribution maps produced by the models were judged for reasonableness based on first-hand knowledge of the species, its habitat preferences, and known range [54]. We further compared model predictions to observational records from Cornell Lab’s eBird citizen-scientist database [55]. We used the area under the operating curve (AUC) [56] as an indication of a model’s classification accuracy (higher values indicating greater accuracy) and the Akaike information criterion corrected for small sample size (AICc) [57] as a measure of relative explanatory power (lower values indicating less information loss). Model similarity was compared with Warren’s I-statistic [58] and Schoener’s D statistic [59] (higher values in both indicating greater similarity) using ENMTools. Single-processor run times were recorded to aid our understanding of algorithm performance and help identify opportunities for multiprocessor parallelization.

## Results

On the basis of permutation importance, 13 of the 19 original bioclim variables were among the top ten most contributory predictors across all three replicated runs of the MaxEnt baseline: bio02, bio03, bio05, bio06, bio08–bio12, bio14, bio15, bio17, and bio18 (Table 2). Of those, bio02, bio05, and bio14 appeared in only one run each at 10th place. Bio18 showed strong dominance throughout. When performance was averaged across all three runs, the top six contributory variables in the ensemble collectively accounted for 65% of overall permutation importance (ensemble average). In descending order of importance, the top six predictors included bio18, bio03, bio10, bio15, bio11, and bio06. When these six top-contributing variables were used in a final MaxEnt run, the model’s four most contributory variables (bio18, bio03, bio10, and bio15) accounted for approximately 86% of overall permutation importance, and its predicted habitat suitability distribution corresponded well with what is known about the natural history of the species and observational records for Cassin’s Sparrow for the year 2016 (Fig 1) [55].

**Fig 1.**
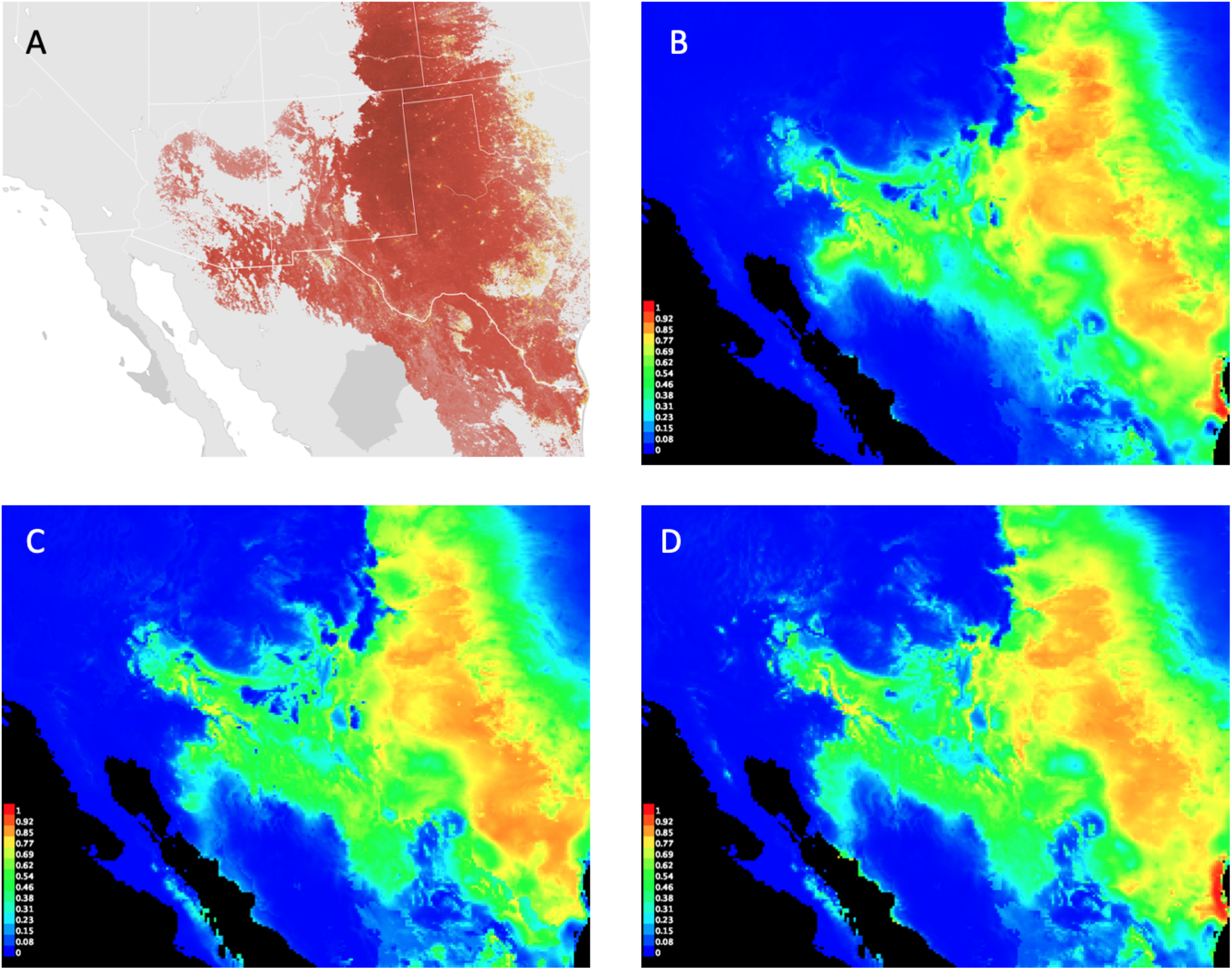
Cassin’s Sparrow distribution maps. Range map depicting the seasonally-averaged estimated relative abundance of Cassin’s Sparrow (A) compared to the species’ predicted habitat suitability distributions obtained from the MaxEnt baseline (B), Monte Carlo Ensemble #1 (C), and Monte Carlo Ensemble #2 (D). Image (A) provided by eBird (www.ebird.org) and created 28 July 2020; images (B)–(D) created by the authors using MaxEnt Version 3.4.1 [61].

**Table 2.**
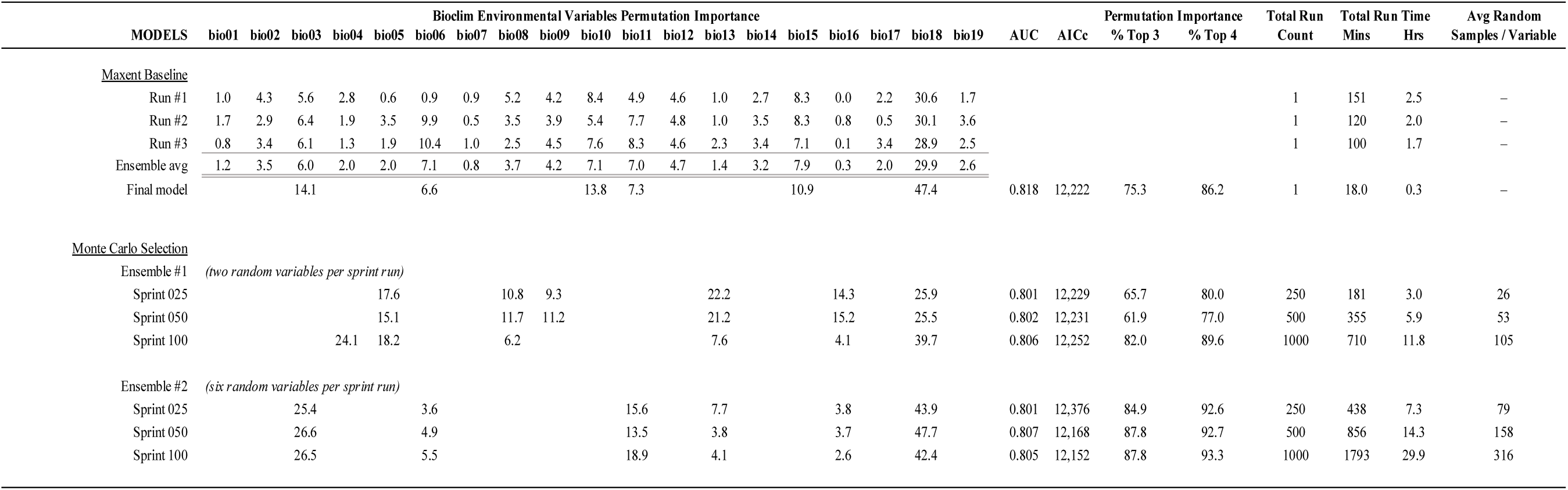
Results of Maxent Baseline and Monte Carlo Selection Trials.

A distinct pattern of progression toward a stable subset of key variables was observed in the Monte Carlo ensembles (Figs 2,3). In both cases, the top three contributory variables among the top six were selected early in the sprint runs, and AICc values fluctuated within a narrow range around an average that changed little over the course of the selection process. Greater variability in the composition of the top six subset was seen in Ensemble #1 where two random variables at a time were selected for each sprint run (Table 2, Fig 2). In Ensemble #2, where six random variables at a time were selected for the MaxEnt runs, the top six variables were identified by the 25th sprint and had settled into their final rank order by sprint 54 (Fig 3). Ensemble #2 appeared to produce the best overall results and shared four variables in common with the top six selected by the MaxEnt baseline (bio03, bio06, bio11, and bio18) (Table 2). Ensemble #2’s final model had the lowest overall AICc, and its four most contributory variables accounted for approximately 93% of overall permutation importance, the highest attained overall.

**Fig 2.**
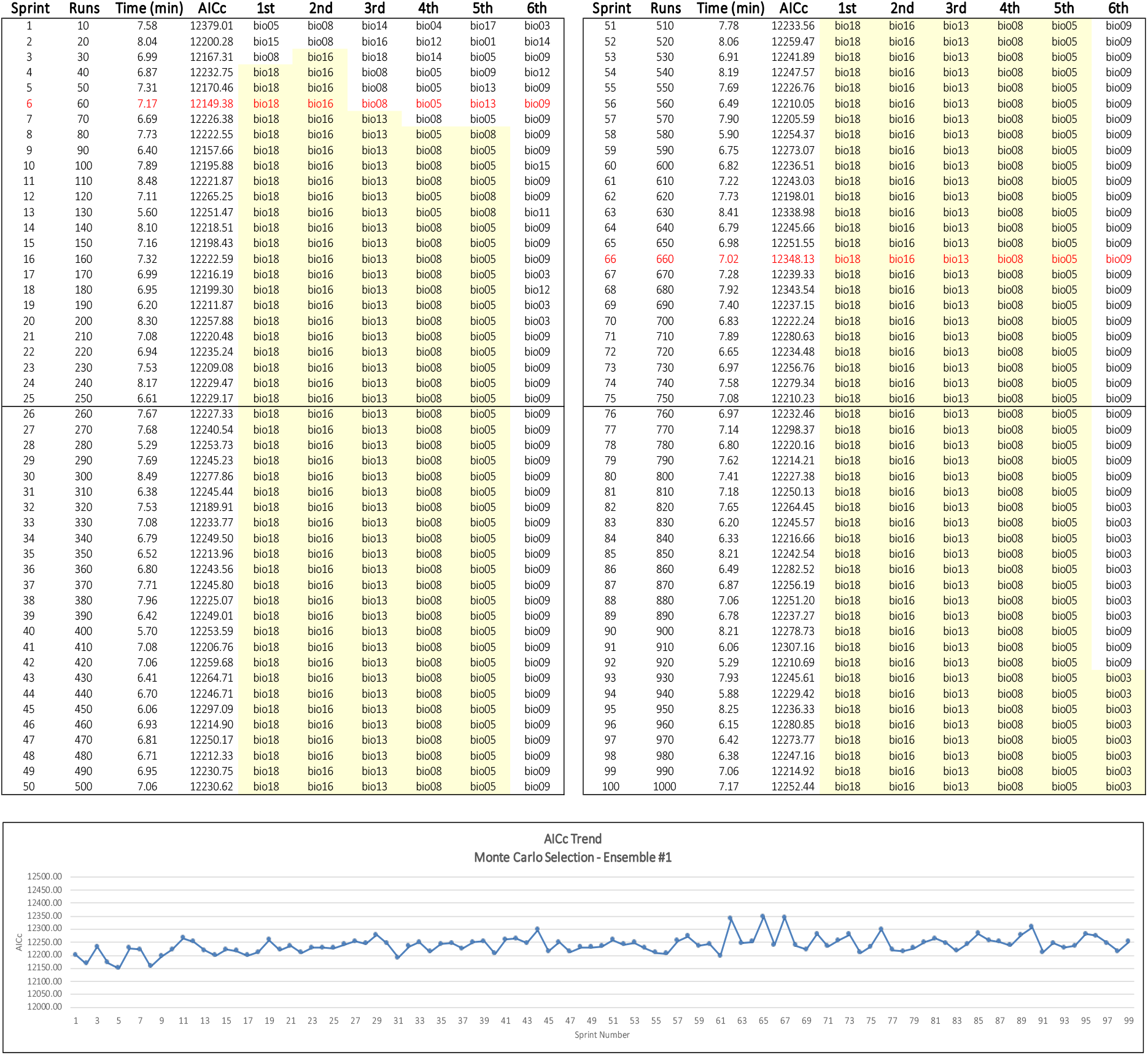
Monte Carlo Ensemble #1 results. Two random variables at a time were chosen for each MaxEnt sprint run. The sprint log on top shows the progressive selection of a stable set of top six variables in yellow. The graph on the bottom shows the narrow range of fluctuating AICc values over the course of the ensemble runs. Maximum and minimum AICc values are shown in red.

**Fig 3.**
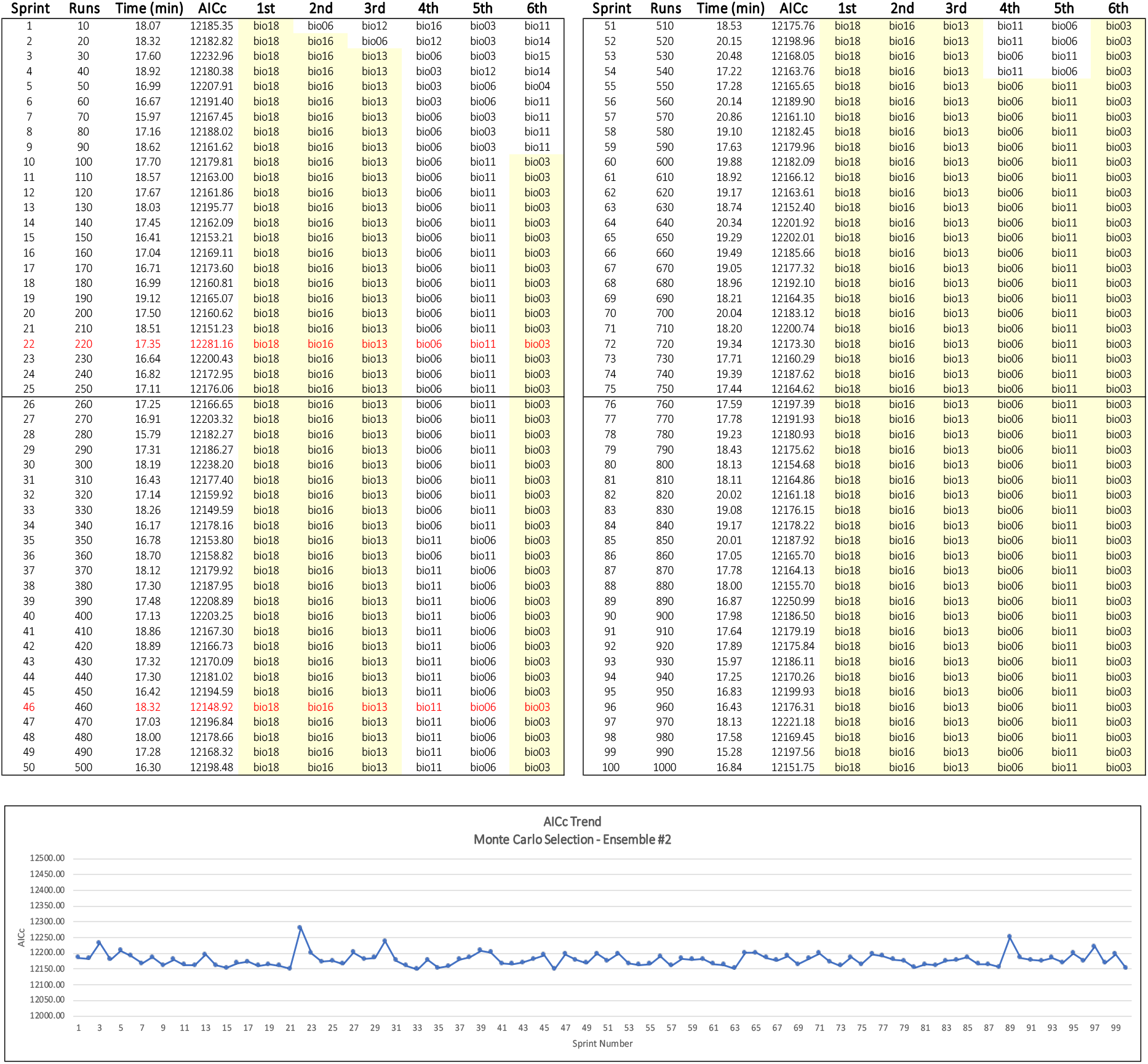
Monte Carlo Ensemble #2 results. Six random variables at a time were chosen for each MaxEnt sprint run. The sprint log on top shows the progressive selection of a stable set of top six variables in yellow. The graph on the bottom shows the narrow range of fluctuating AICc values over the course of the ensemble runs. Maximum and minimum AICc values are shown in red.

Ensemble #1 had only one variable in common with the top six selected by both the baseline run and Ensemble #2. What accounts for this difference is not immediately apparent; however, we speculate that the random pair-wise comparisons occurring in Ensemble #1 may alter the relative global influence of the collinearities known to exist in the bioclim variables [62–64]. The average number of times a variable was sampled appeared to have a marginal, positive influence on resulting model quality once an adequate minimum was attained. Ensemble #2 results suggest that at least 80 uniformly distributed samples per starting-set variable are needed to identify a reasonable top six set of variables; the best overall model resulted from over 300 samples per variable (Table 2).

## Discussion

The most striking outcome of the study is the similarity in results. Final models in the MaxEnt baseline and the Monte Carlo ensembles produced predicted habitat suitability distributions that are nearly indistinguishable from one another (Fig 1). Internal metrics likewise reveal little difference in outcomes, with AUC values ranging only from 0.801 to 0.818 and AICc ranging from 12,152 to 12,222, suggesting that the traditional MaxEnt runs and the Monte Carlo approach both produced reasonable models (Table 2). The two approaches also each identified four variables that collectively contributed more than 80% to the formulation of their respective models. Across the board, models showed a high degree of similarity in Schoener’s D and the I-statistic (Table 3).

**Table 3.** Model Similarity Metrics.

| <u>Schoener's D Statistic</u> |  |  |  |  |  |  |  |  |  |  |
| --- | --- | --- | --- | --- | --- | --- | --- | --- | --- | --- |
| MODELS ↓ → | Maxent-Run1 | Maxent-Run2 | Maxent-Run3 | Maxent-Final | MC-E1-025 | MC-E1-050 | MC-E1-100 | MC-E2-025 | MC-E2-050 | MC-E2-100 |
| Maxent-Run1 | 1 | 0.9712 | 0.9738 | 0.9355 | 0.8629 | 0.8630 | 0.9044 | 0.8882 | 0.8903 | 0.8906 |
| Maxent-Run2 | x | 1 | 0.9793 | 0.9397 | 0.8639 | 0.8648 | 0.9078 | 0.8915 | 0.8949 | 0.8947 |
| Maxent-Run3 | x | x | 1 | 0.9354 | 0.8579 | 0.8586 | 0.9039 | 0.8871 | 0.8903 | 0.8902 |
| Maxent-Final | x | x | x | 1 | 0.8667 | 0.8673 | 0.9214 | 0.9104 | 0.9138 | 0.9142 |
| MC-E1-025 | x | x | x | x | 1 | 0.9880 | 0.9006 | 0.8810 | 0.8801 | 0.8785 |
| MC-E1-050 | x | x | x | x | x | 1 | 0.9026 | 0.8813 | 0.8804 | 0.8786 |
| MC-E1-100 | x | x | x | x | x | x | 1 | 0.9393 | 0.9389 | 0.9365 |
| MC-E2-025 | x | x | x | x | x | x | x | 1 | 0.9815 | 0.9810 |
| MC-E2-050 | x | x | x | x | x | x | x | x | 1 | 0.9844 |
| MC-E2-100 | x | x | x | x | x | x | x | x | x | 1 |
| <u>Warren's I Statistic</u> |  |  |  |  |  |  |  |  |  |  |
| MODELS ↓ → | Maxent-Run1 | Maxent-Run2 | Maxent-Run3 | Maxent-Final | MC-E1-025 | MC-E1-050 | MC-E1-100 | MC-E2-050 | MC-E2-100 | MC-E2-100 |
| Maxent-Run1 | 1 | 0.9991 | 0.9994 | 0.9948 | 0.9760 | 0.9758 | 0.9874 | 0.9828 | 0.9833 | 0.9834 |
| Maxent-Run2 | x | 1 | 0.9995 | 0.9949 | 0.9760 | 0.9760 | 0.9878 | 0.9831 | 0.9838 | 0.9839 |
| Maxent-Run3 | x | x | 1 | 0.9945 | 0.9748 | 0.9748 | 0.9869 | 0.9823 | 0.9830 | 0.9831 |
| Maxent-Final | x | x | x | 1 | 0.9755 | 0.9753 | 0.9894 | 0.9871 | 0.9878 | 0.9880 |
| MC-E1-025 | x | x | x | x | 1 | 0.9998 | 0.9887 | 0.9798 | 0.9797 | 0.9784 |
| MC-E1-050 | x | x | x | x | x | 1 | 0.9889 | 0.9795 | 0.9794 | 0.9781 |
| MC-E1-100 | x | x | x | x | x | x | 1 | 0.9910 | 0.9912 | 0.9906 |
| MC-E2-025 | x | x | x | x | x | x | x | 1 | 0.9996 | 0.9994 |
| MC-E2-050 | x | x | x | x | x | x | x | x | 1 | 0.9996 |
| MC-E2-100 | x | x | x | x | x | x | x | x | x | 1 |

The most significant drawback identified in the study was the long run times. MaxEnt’s linear scaling behavior can be challenging in a single-processor environment. In the baseline runs, producing a single model through MaxEnt’s GUI using our selected settings involved writing many files to disk and took from 18 minutes (with six variables) to over two hours (with all 19 variables). MaxEnt in the R environment outputs memory-resident objects, which results in faster run times. Still, with its repeated invocations of MaxEnt, Ensemble #2 took nearly 30 hours to complete (Table 2).

The Monte Carlo approach also exhibits linear scaling properties; however, its scaling behavior is not determined by the MaxEnt program, since each of the MaxEnt runs in the Monte Carlo method operates on a set number of predictors. The method’s linear scaling property is determined, instead, by the need to adequately sample the starting set of environmental variables in order to obtain a good result. What makes a practical implementation of the Monte Carlo method possible is that each of its MaxEnt runs is entirely independent from all other runs in the ensemble. This high level of subtask independence is sometimes referred to as an “embarrassingly parallel” workload, which makes it relatively straightforward to implement in a cluster computing environment. If 1000 processors were recruited into service — which is becoming increasingly convenient with the proliferation of multiprocessor, high performance cloud computing — a 1000-run ensemble could conceivably take about as long as a single MaxEnt run.

The potential significance of this advantage becomes apparent when one considers the method’s use with large collections of environmental data. The Monte Carlo approach described here provides an approximate solution to the problem of finding a useful *k*-size subset of an *n*-size collection of variables. In principal, there are *n*! / [*k*!(*n-k*)!] variable combinations to consider in such an evaluation, a staggering 27,000-plus six-variable subsets with the 19 bioclim variables alone. Algorithms that accomplish variable selection through stepwise removal or are otherwise bound to the linear scaling properties of underlying software components are inherently unable to exhaustively explore this combinatorial space. A Monte Carlo method makes such a search possible by randomly sampling the universe of possible combinations and returning approximate solutions in practical amounts of time, particularly if implemented as a high-performance cloud service.

The use of IPCC-class climate model outputs in efforts to assess the impacts of climate change on biodiversity and other ecosystem processes is growing. Exploring the potential of these massive data sets, expanded use of ensemble modeling, and the actual work of fitting models for the thousands of species scientists wish to study will require hundreds to thousands of projections [16,23]. An improved capacity to use large environmental data sets in MaxEnt modeling would benefit this work. We are encouraged to think that innovative use of Monte Carlo techniques might provide a helpful means of meeting this challenge.

## Conclusions

This small-scale, proof-of-concept study leaves many practical and theoretical questions unanswered. Preliminary results, however, suggest that a Monte Carlo method could offer a viable means of selecting environmental predictors for MaxEnt models that is amenable to parallelization and scalable to large data sets, including externally-stored collections. This points to the possibility of near-real-time multiprocessor implementations that could enable broader and more exploratory use of global climate model outputs in environmental niche modeling and aid in the discovery of viable predictors. Next steps will focus on implementing this capability in NASA’s Advanced Data Analytics Platform (ADAPT) science cloud, evaluating the method’s behavior using products generated by the Goddard Earth Observing System, Version 5 (GEOS-5) modeling system, extending stochasticity to feature class and regularization multiplier selection, developing automatic stopping rules, and evaluating the method’s effectiveness in addressing research questions relating to climate change influences on Cassin’s Sparrow abundance and distribution.

## Author contributions

JLS: Conceptualization, Formal analysis, Investigation, Methodology, Writing – original draft. MLC: Conceptualization, Formal analysis, Methodology, Writing – review & editing. RLG: Software. GST: Software. JL: Software. SLS: Validation. TPM: Software. MEA: Visualization.

## References

1. Elith J, Phillips SJ, Hastie T, Dudík M, Chee YE, Yates CJ. A statistical explanation of MaxEnt for ecologists. Diversity and distributions. 2011;17: 43–57.

2. Phillips SJ, Anderson RP, Dudík M, Schapire RE, Blair ME. Opening the black box: An open-source release of Maxent. Ecography. 2017;40: 887–893.

3. Phillips SJ. A brief tutorial on Maxent. AT&T Research. 2005;190: 231–259.

4. Phillips SJ, Anderson RP, Schapire RE. Maximum Entropy Modeling of Species Geographic Distributions. Ecological Modelling. 2006;190: 231–259. doi:10.1016/j.ecolmodel.2005.03.026.

5. Kalinski CE. Building Better Species Distribution Models with Machine Learning: Assessing the Role of Covariate Scale and Tuning in Maxent Models. 2019; 129.

6. Merow C, Smith MJ, Silander JA. A practical guide to MaxEnt for modeling species’ distributions: what it does, and why inputs and settings matter. Ecography. 2013;36: 1058–1069. doi:10.1111/j.1600-0587.2013.07872.x

7. Schmidt-Lebuhn AN, Knerr NJ, Miller JT, Mishler BD. Phylogenetic diversity and endemism of Australian daisies (Asteraceae). Journal of Biogeography. 2015;42: 1114–1122. doi:10.1111/jbi.12488

8. Warren DL, Wright AN, Seifert SN, Shaffer HB. Incorporating model complexity and spatial sampling bias into ecological niche models of climate change risks faced by 90 California vertebrate species of concern. Diversity and distributions. 2014;20: 334–343.

9. Cardoso-Leite R, Vilarinho AC, Novaes MC, Tonetto AF, Vilardi GC, Guillermo-Ferreira R. Recent and future environmental suitability to dengue fever in Brazil using species distribution model. Transactions of The Royal Society of Tropical Medicine and Hygiene. 2014;108: 99–104. doi:10.1093/trstmh/trt115

10. Morisette JT, Jarnevich CS, Ullah A, Cai W, Pedelty JA, Gentle JE, et al. A Tamarisk Habitat Suitability Map for the Continental United States. Frontiers in Ecology and the Environment. 2006;4: 11–17. doi:10.1890/1540-9295(2006)004

11. Stohlgren TJ, Schnase JL. Risk Analysis for Biological Hazards: What We Need to Know about Invasive Species. Risk Analysis. 2006;26: 163–73. doi:10.1111/j.1539-6924.2006.00707.x.

12. Beauchamp VB, Koontz SM, Suss C, Hawkins C, Kyde KL, Schnase JL. An Introduction to Oplismenus Undulatifolius (Ard.) Roem. & Schult (Wavyleaf Basketgrass), a Recent Invader in Mid-Atlantic Forest Understories 1,2. The Journal of the Torrey Botanical Society. 2013;140: 391–413. doi:10.3159/torrey-d-13-00033.1.

13. alwi Muttaqin luthfi, Heru Murti S, Susilo B. MaxEnt (Maximum Entropy) model for predicting prehistoric cave sites in Karst area of Gunung Sewu, Gunung Kidul, Yogyakarta. In: Wibowo SB, Rimba AB, A. Aziz A, Phinn S, Sri Sumantyo JT, Widyasamratri H, et al., editors. Sixth Geoinformation Science Symposium. Yogyakarta, Indonesia: SPIE; 2019. p. 3. doi:10.1117/12.2543522

14. Feng X, Park DS, Walker C, Peterson AT, Merow C, Papeş M. A checklist for maximizing reproducibility of ecological niche models. Nature Ecology & Evolution. 2019;3: 1382–1395. doi:10.1038/s41559-019-0972-5

15. Morales NS, Fernández IC, Baca-González V. MaxEnt’s parameter configuration and small samples: are we paying attention to recommendations? A systematic review. PeerJ. 2017;5: e3093. doi:10.7717/peerj.3093

16. Araújo M, New M. Ensemble forecasting of species distributions. Trends in Ecology & Evolution. 2007;22: 42–47. doi:10.1016/j.tree.2006.09.010

17. Zeng Y, Low BW, Yeo DCJ. Novel methods to select environmental variables in MaxEnt: A case study using invasive crayfish. Ecological Modelling. 2016;341: 5–13. doi:https://doi.org/10.1016/j.ecolmodel.2016.09.019

18. Araújo MB, Anderson RP, Barbosa AM, Beale CM, Dormann CF, Early R, et al. Standards for distribution models in biodiversity assessments. Science Advances. 2019;5: eaat4858.

19. Qiao H, Soberón J, Peterson AT. No silver bullets in correlative ecological niche modelling: insights from testing among many potential algorithms for niche estimation. Kriticos D, editor. Methods in Ecology and Evolution. 2015;6: 1126–1136. doi:10.1111/2041-210X.12397

20. Ashraf U, Peterson AT, Chaudhry MN, Ashraf I, Saqib Z, Rashid Ahmad S, et al. Ecological niche model comparison under different climate scenarios: a case study of *Olea* spp. in Asia. Ecosphere. 2017;8: e01825. doi:10.1002/ecs2.1825

21. Guisan A, Zimmermann NE. Predictive habitat distribution models in ecology. Ecological modelling. 2000;135: 147–186.

22. Li Y, Li M, Li C, Liu Z. Optimized Maxent Model Predictions of Climate Change Impacts on the Suitable Distribution of Cunninghamia lanceolata in China. Forests. 2020;11: 302. doi:10.3390/f11030302

23. Cavanagh RD, Murphy EJ, Bracegirdle TJ, Turner J, Knowland CA, Corney SP, et al. A Synergistic Approach for Evaluating Climate Model Output for Ecological Applications. Frontiers in Marine Science. 2017;4: 308. doi:10.3389/fmars.2017.00308

24. Harris RMB, Grose MR, Lee G, Bindoff NL, Porfirio LL, Fox-Hughes P. Climate projections for ecologists: Climate projections for ecologists. Wiley Interdisciplinary Reviews: Climate Change. 2014;5: 621–637. doi:10.1002/wcc.291

25. Stock CA, Alexander MA, Bond NA, Brander KM, Cheung WW, Curchitser EN, et al. On the use of IPCC-class models to assess the impact of climate on living marine resources. Progress in Oceanography. 2011;88: 1–27.

26. Bojinski S, Verstraete M, Peterson TC, Richter C, Simmons A, Zemp M. The Concept of Essential Climate Variables in Support of Climate Research, Applications, and Policy. Bulletin of the American Meteorological Society. 2014;95: 1431–1443. doi:10.1175/BAMS-D-13-00047.1

27. Braunisch V, Coppes J, Arlettaz R, Suchant R, Schmid H, Bollmann K. Selecting from correlated climate variables: a major source of uncertainty for predicting species distributions under climate change. Ecography. 2013;36: 971–983. doi:10.1111/j.1600-0587.2013.00138.x

28. Schnase JL. Climate Analytics as a Service. Cloud Computing in Ocean and Atmospheric Sciences. 2016. pp. 187–219. doi:10.1016/b978-0-12-803192-6.00011-6

29. Edwards PN. A Vast Machine: Computer Models, Climate Data, and the Politics of Global Warming. Cambridge, MA: MIT Press; 2010.

30. IPCC — Intergovernmental Panel on Climate Change. 2020 [cited 14 Mar 2020]. Available: https://www.ipcc.ch/

31. Responding to the Challenge of Climate and Environmental Change: NASA’s Plan for a Climate-Centric Architecture for Earth Observations and Applications from Space. National Aeronautics and Space Administration; 2010. Available: https://gmao.gsfc.nasa.gov/reanalysis/MERRA/.

32. Araújo MB, Guisan A. Five (or so) challenges for species distribution modelling. Journal of Biogeography. 2006;33: 1677–1688. doi:10.1111/j.1365-2699.2006.01584.x

33. Duan Y, Edwards JS, Dwivedi YK. Artificial intelligence for decision making in the era of Big Data – evolution, challenges and research agenda. International Journal of Information Management. 2019;48: 63–71. doi:10.1016/j.ijinfomgt.2019.01.021

34. Galante PJ, Alade B, Muscarella R, Jansa SA, Goodman SM, Anderson RP. The challenge of modeling niches and distributions for data-poor species: a comprehensive approach to model complexity. Ecography. 2018;41: 726–736.

35. Muscarella R, Galante PJ, Soley-Guardia M, Boria RA, Kass JM, Uriarte M, et al. ENMeval: An R package for conducting spatially independent evaluations and estimating optimal model complexity for Maxent ecological niche models. Methods in Ecology and Evolution. 2014;5: 1198–1205. doi:10.1111/2041-210X.12261

36. Radosavljevic A, Anderson RP. Making better MaxEnt models of species distributions: complexity, overfitting and evaluation. Araújo M, editor. Journal of Biogeography. 2014;41: 629–643. doi:10.1111/jbi.12227

37. Kroese DP, Brereton T, Taimre T, Botev ZI. Why the Monte Carlo method is so important today: Why the MCM is so important today. Wiley Interdisciplinary Reviews: Computational Statistics. 2014;6: 386–392. doi:10.1002/wics.1314

38. Dunning, Jr. JB, Bowers, Jr. RK, Suter SJ, Bock CE. Cassin’s Sparrow (Peucaea cassinii), Version 1.0. In: Birds of the World (P. G. Rodewald, Editor) [Internet]. 2020 [cited 22 May 2020]. Available: https://doi.org/10.2173/bow.casspa.01

39. Iknayan KJ, Beissinger SR. Collapse of a desert bird community over the past century driven by climate change. Proc Natl Acad Sci USA. 2018;115: 8597. doi:10.1073/pnas.1805123115

40. Radchuk V, Reed T, Teplitsky C, van de Pol M, Charmantier A, Hassall C, et al. Adaptive responses of animals to climate change are most likely insufficient. Nature Communications. 2019;10: 3109. doi:10.1038/s41467-019-10924-4

41. GBIF.org (21 February 2019) GBIF Occurrence Download https://doi.org/10.15468/dl.0s8yak.

42. Fick SE, Hijmans RJ. WorldClim 2: new 1-km spatial resolution climate surfaces for global land areas. International Journal of Climatology. 2017;37: 4302–4315. doi:10.1002/joc.5086

43. Worldclim bioclimatic variables. 2020 [cited 22 May 2020]. Available: https://worldclim.org/data/worldclim21.html

44. GDAL/OGR Geospatial Data Abstraction Software Library. Open Source Geospatial Foundation; 2020. Available: https://gdal.org/

45. Hijmans RJ, Phillips S, Elith J, Leathwick J. dismo: Species Distribution Modeling. 2017. Available: https://CRAN.R-project.org/package=dismo

46. Feng X, Park DS, Liang Y, Pandey R, Papeş M. Collinearity in ecological niche modeling: Confusions and challenges. Ecology and Evolution. 2019;9: 10365–10376. doi:10.1002/ece3.5555

47. Maxent Version 3.4.1 Download Site. [cited 22 May 2020]. Available: https://biodiversityinformatics.amnh.org/open_source/maxent/

48. R: The R Project for Statistical Computing. [cited 22 May 2020]. Available: https://www.r-project.org/

49. Muscarella R, Galante PJ, Soley-Guardia M, Boria RA, Kass JM, Anderson MU and RP. ENMeval: Automated Runs and Evaluations of Ecological Niche Models. 2018. Available: https://CRAN.R-project.org/package=ENMeval

50. RStudio | Open source & professional software for data science teams. [cited 27 May 2020]. Available: https://rstudio.com/

51. Warren DL, Glor RE, Turelli M. ENMTools: a toolbox for comparative studies of environmental niche models. Ecography. 2010 [cited 27 Mar 2020]. doi:10.1111/j.1600-0587.2009.06142.x

52. Phillips SJ, Dudík M, Elith J, Graham CH, Lehmann A, Leathwick J, et al. Sample selection bias and presence-only distribution models: implications for background and pseudo-absence data. Ecological applications. 2009;19: 181–197.

53. Fourcade Y, Engler JO, Rödder D, Secondi J. Mapping Species Distributions with MAXENT Using a Geographically Biased Sample of Presence Data: A Performance Assessment of Methods for Correcting Sampling Bias. Valentine JF, editor. PLoS ONE. 2014;9: e97122. doi:10.1371/journal.pone.0097122

54. Schnase JL, Grant WE, Maxwell TC, Leggett JJ. Time and energy budgets of Cassin’s sparrow (Aimophila cassinii) during the breeding season: evaluation through modelling. Ecological Modelling. 1991;55: 285–319.

55. (Peucaea cassinii) - Species Map - eBird. 2020 [cited 31 May 2020]. Available: https://ebird.org/map/casspa

56. Fielding AH, Bell JF. A review of methods for the assessment of prediction errors in conservation presence/absence models. Environmental Conservation. 1997;24: 38–49. doi:10.1017/S0376892997000088

57. H. Akaike. A new look at the statistical model identification. IEEE Transactions on Automatic Control. 1974;19: 716–723. doi:10.1109/TAC.1974.1100705

58. Warren DL, Glor RE, Turelli M. Environmental niche equivalency versus conservatism: quantitative approaches to niche evolution. Evolution. 2008;62: 2868–2883. doi:10.1111/j.1558-5646.2008.00482.x

59. Schoener TW. The Anolis Lizards of Bimini: Resource Partitioning in a Complex Fauna. Ecology. 1968;49: 704–726. doi:10.2307/1935534

60. Fink D, Auer T, Johnston A, Strimas-Mackey M, Robinson O, Ligocki S, et al. Cassin’s Sparrow - Abundance map - eBird Status and Trends. In: eBird Status and Trends, Data Version: 2018; Released: 2020 [Internet]. 2020 [cited 5 Oct 2020]. Available: https://ebird.org/ebird/science/status-and-trends/casspa/abundance-map

61. Phillips SJ, Dudík M, Schapire RE. Maxent software for modeling species niches and distributions (Version 3.4.1). 2020. Available: url: http://biodiversityinformatics.amnh.org/open_source/maxent/

62. Tang Y, Winkler JA, Viña A, Liu J, Zhang Y, Zhang X, et al. Pearson pairwise correlation matrix between the bioclimatic variables. 2018. doi: 10.1371/journal.pone.0189496.g003

63. O’Donnell MS, Ignizio D a. Bioclimatic Predictors for Supporting Ecological Applications in the Conterminous United States. Reston, VA: US Geological Survey; 2012 p. 10. Report No.: 691. Available: https://pubs.usgs.gov/ds/691/

64. Dormann CF, Elith J, Bacher S, Buchmann C, Carl G, Carré G, et al. Collinearity: a review of methods to deal with it and a simulation study evaluating their performance. Ecography. 2013;36: 27–46. doi:10.1111/j.1600-0587.2012.07348.x

